# Inhibition of (interstitial) P2Y_6_ receptors attenuates renal fibrosis progression

**DOI:** 10.64898/2026.01.27.701928

**Authors:** Lena Marie Süß, Anna Petzendorfer, Bettina Firmke, Anja Süß, Richard Warth, Katharina Anna-Elisabeth Broeker, Anna-Lena Forst

**Author notes:** Corresponding author: Dr. rer. nat. Anna-Lena Forst, Medical Cell Biology, University of Regensburg, Universitätsstr. 31, D-93053 Regensburg, Germany.

## Abstract

Chronic kidney disease (CKD) affects over 850 million people worldwide and is characterized by progressive renal fibrosis driven by activated interstitial fibroblasts. Signaling by extracellular nucleotides and P2 receptors plays an important role in renal pathophysiology, yet its contribution to fibroblast activation and fibrosis remains poorly understood. Here, we investigated the expression and function of G_q/11_-coupled P2Y receptors in renal interstitial fibroblasts and their involvement in experimental kidney fibrosis.

Using highly selective RNA in situ hybridization, we detected P2Y_1_ (*P2ry1*) and P2Y_6_ (*P2ry6*) receptor expression in interstitial fibroblasts. Notably, P2Y_6_ expression was markedly upregulated in several experimental mouse models of renal fibrosis. Functional assays in primary cultured renal fibroblasts confirmed G_q/11_-coupled P2Y receptor activity, as evidenced by transient intracellular Ca^2+^ elevations upon nucleotide stimulation. Primary cultured renal fibroblasts exhibited enhanced migration in response to extracellular uridine diphosphate (UDP). To assess the contribution of interstitial P2Y_6_ receptors to fibrosis progression, we employed an adenine-induced nephropathy model with or without the selective P2Y_6_ antagonist MRS2578. Pharmacological inhibition of P2Y_6_ significantly reduced the mRNA expression of the myofibroblast marker α-smooth muscle actin and collagen I.

Collectively, these findings suggest that upregulated P2Y_6_ receptor signaling promotes the transition of resident interstitial cells into myofibroblasts during renal fibrosis, likely by modulating fibroblast migration. Inhibition of P2Y6 signaling could represent a new strategy for reducing excessive renal fibrosis.

**TRANSLATIONAL STATEMENT:** This study reveals the role of the P2Y_6_ receptor (*P2ry6*) in fibrotic processes in the kidney. P2Y_6_, a G_q/11_ protein-coupled UDP-sensitive receptor, is expressed in renal interstitial PDGFR-β-positive cells and macrophages. Its pharmacological inhibition significantly reduces fibrosis in the mouse adenine nephropathy model. Blocking P2Y_6_ therefore represents a promising therapeutic strategy for kidney diseases characterized by excessive scarring.

## INTRODUCTION

Renal fibrosis represents a common pathological end-stage of chronic kidney disease (CKD) and is a major contributor to progressive loss of kidney function. It is characterized by excessive accumulation of extracellular matrix (ECM) components such as collagens, fibronectin, and tenascins, leading to structural alterations and ultimately irreversible damage to the renal parenchyma and loss of endocrine functions ^1–3^. Increased ECM production primarily originates from myofibroblasts, which can transdifferentiate from several cell types, including fibroblasts, monocytes, tubular epithelial cells, and endothelial cells ^4–7^. Although the relative contribution of these cell types to the myofibroblast population may vary depending on the experimental model, growing evidence identifies platelet-derived growth factor receptor-β (PDGFR-β)-expressing resident fibroblasts and pericytes as the main precursors of myofibroblasts in human kidneys ^2,8,9^. Macrophage-to-myofibroblast transition also contributes to interstitial fibrosis in chronic renal allograft injury ^10,11^.

Differentiation of PDGFR-β positive cells into myofibroblasts is a multifactorial process involving numerous signaling pathways, including the well-studied transforming growth factor-β1 (TGF-β1) and angiotensin II ^3,12,13^. In addition to these established profibrotic mediators, recent studies implicate extracellular nucleotides as important regulators of fibroblast homeostasis. Nucleotide release is likely regulated and facilitated by several mechanisms, including vesicular or lysosomal exocytosis and channel-mediated release via connexins or pannexins during cellular stress such as mechanical strain or hypoxia. These nucleotides act as danger-associated molecular patterns (DAMPs) in both paracrine and autocrine signaling ^14–17^. Extracellular nucleotides act through specific P2 and P1 receptors ^15,18–21^. While P1 receptors bind adenosine, P2 receptors are activated by purine and pyrimidine nucleotides and can be further subdivided into P2Y receptors and P2X receptors. P2X receptors function as non-selective cation channels, while the P2Y receptor isoforms are G-protein coupled receptors that are activated with differing selectivity by adenosine triphosphate (ATP), adenosine diphosphate (ADP), uridine triphosphate (UTP), and uridine diphosphate (UDP) ^22,23^. P1 and P2 receptor expression has been reported in all segments of the nephron and renal vasculature. Epithelial cells often express multiple receptor subtypes at both the apical and basolateral cell membranes ^15,19^, yet little is known about expression of these receptors in interstitial cells.

Besides P1 and P2 receptors, other components of the purinergic signaling pathway include ectonucleases like CD39 (*ENTPD1*), which catalyze sequential hydrolysis of tri- and diphosphates to monophosphates or CD73 (*NT5E*) converting adenosine monophosphate to adenosine. Notably, CD73 is another marker for interstitial fibroblasts that is expressed by appr. 60% of PDGFR-β-positive cells indicative of active purinergic activity in the proximity of interstitial cells ^8^. It is well known that important physiological functions such as cell proliferation and growth, energy metabolism, and transepithelial flow are influenced by extracellular nucleotides. However, knowledge about the significance of these signaling pathways in interstitial cells of the kidney is still incomplete.

In the pathophysiological setting, renal P2R receptor activation occurs in diverse inflammatory and non-inflammatory diseases including hypertension ^19,24–26^, transplant rejection ^21^ and polycystic kidney disease ^20,27^. The influence of P1 or P2 receptors on the development of renal fibrosis has been the subject of several studies with context-depending and sometimes controversial results: Adenosine signaling through P1 receptors, particularly A2A and A2B, exerted protective as well as profibrotic effects ^21^. Abrogating P2X_7_ receptor signaling, which is known to promote inflammation and fibrotic remodeling via NLRP3 inflammasome activation and IL-1β release, promised the most therapeutic potential in murine fibrotic disease models so far ^24,25,28,29^. Despite these promising pre-clinical results however, the beneficial effect in completed phase 2 clinical trials was disappointingly mild ^15^.

While the current research focuses on the inhibition of inflammatory cells via blockage of P2X_7_, we were interested in the potential role of P2Y receptors of interstitial cells in fibrosis progression. In the present study, we analyzed the expression and functionality of murine G_q/11_-protein coupled receptors (P2Y_1_, P2Y_2_, P2Y_4_ and P2Y_6_) in healthy and fibrotic murine kidneys. Additionally, we subjected mice to different experimental models of kidney fibrosis to elucidate the therapeutic potential of specific P2Y_6_ signaling on fibrosis progression.

## METHODS

### Ethical Approval

All animal experiments were conducted in accordance with Directive 2010/63/EU of the European Parliament and of the Council on the protection of animals used for scientific purposes. The experiments also comply with the animal ethics checklist of this journal (Grundy, 2015). All experiments were approved the local councils for animal care (Regierung von Unterfranken) according to the German law for animal care.

### Mice

Wildtype mice in a BL/6J background were used in the studies with experimental kidney fibrosis. For FACS-sorting of PDGFR-β-positive renal cells, murine kidneys from tamoxifen-induced PDGFR-β Cre^ERT/2^ mTmG mice ^30^ that express a membranous GFP under control of the PDGFR-β-promotor were used. All mice were kept on 12∶12 hour light-dark cycle, controlled temperature levels (22°C ± 2°C) and humidity (55 % ± 10 %). Animals were fed a standard rodent chow (0.6% NaCl; Ssniff, Soest, Germany) with free access to autoclaved tap water.

### Adenine-induced nephropathy

Adenine-induced fibrosis was generated in adult male mice at the age of 6-16 weeks. An 0.2% adenine-containing diet (altromin Spezialfutter GmbH, Germany) was fed continuously for 3 weeks. Experiments were performed after exactly 3 weeks (3-week adenine).

### (Reversable) unilateral ureteral obstruction (UUO)

For analysis of UUO kidneys, paraffin-embedded tissue samples from Fuchs and colleagues were re-examined, thereby avoiding additional animal experimentation and ensuring that the procedure is fully compliant with the 3R guidelines for animal research ^3^. In short, ureteral ligation using a suture was performed close to the right kidney through a small abdominal incision under inhalation anesthesia. Five days after the procedure, mice were killed and perfused for RNAscope or kidneys were removed for mRNA quantification. For reversable unilateral ureteral obstruction (rUUO) kidneys, the right kidney of male mice was clipped at the age of 6-16 weeks by making a small incision in the abdomen under anesthesia using 0.5 mg*kg^-1^ medetomidine, 5 mg/kg, midazolam, and 0.05 mg*kg^-1^, fentanyl at day 0. In addition, 200 mg*kg^-1^ paracetamol and 25 mg*kg^-1^ tramadol was administered subcutaneously for pain relief. To awaken the mice from anesthesia, they were injected subcutaneously with 2.5 mg*kg^-1^ atipamezole, 0.5 mg*kg^-1^ flumazenil, and 1.2 mg*kg^-1^ naloxone. The clip was replaced more caudally at day 2 and removed at day 4, whereafter mice were left to recover for three weeks.

### Drug Treatments

For *in vivo* use, MRS2578 (Selleckchem Chemicals LLC, Houston, USA) was prepared at a concentration of 5 mg*mL^-1^ in a vehicle consisting of 30% propylene glycol, 5% Tween 80 and 65% D5W according to the manufacturer’s instructions and administered i.p. at a concentration of 10 mg*kg^-1^ BW three times a week. Treatment started one day before start of the experiment while control animals received solvent-injections intraperitoneally (2 µl*kg^-1^ BW) at the same time points.

For *in vitro* use, a stock solution of 5 mM MRS2578 was prepared in DMSO and used at 5 µM concentration.

### Determination of mRNA expression by real-time PCR

Total RNA was isolated from murine kidneys after perfusion with 0.9% NaCl containing heparin. Kidneys were snap frozen in liquid nitrogen, total RNA was extracted using the RNeasy plus mini kit (Quiagen, Hilden, Germany). The purity and integrity of the RNA were verified spectroscopically using a Nano Drop spectrometer (Life Technologies GmbH, Darmstadt, Germany). For qPCR, cDNA was generated from 1 µg total RNA by reverse transcription using M-MLV Reverse Transcriptase (Life Technologies GmbH, Darmstadt, Germany) according to the protocol provided. To quantify mRNA expression, real-time PCR was performed using the LightCycler Takyon® No ROX SYBR 2X MasterMix (Eurogentec, Seraing, Belgium) and the LightCycler 96 SW instrument (Roche Diagnostics, Mannheim, Germany). Transcript levels were normalized to the expression of the housekeeping protein β-Actin (*Actb*). Primers (Eurofins, Munich, Germany) are listed in table 1.

**Table 1:**
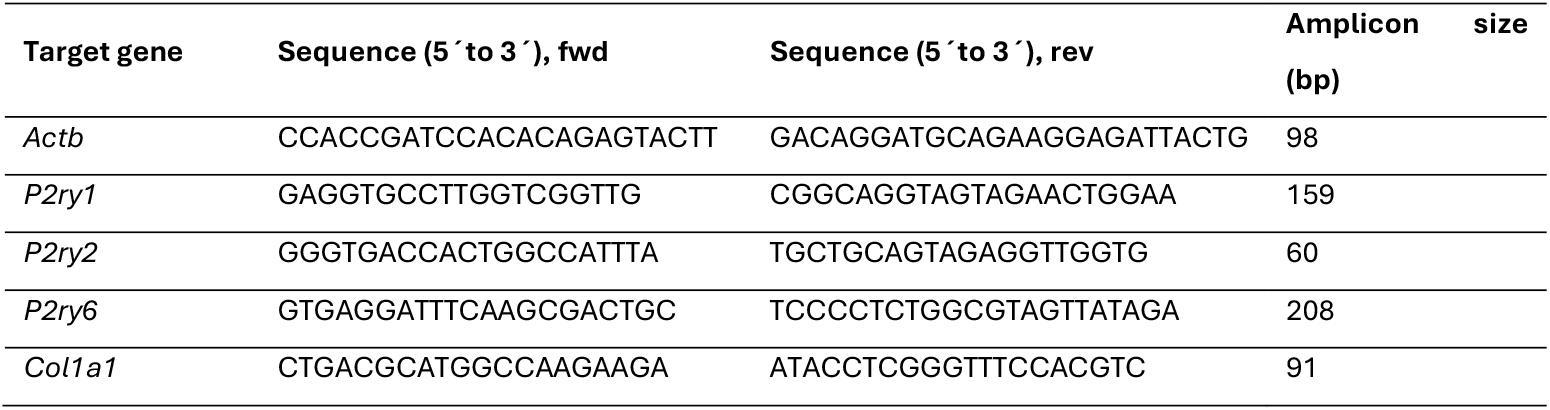
Primer sequences used for qPCR.

### In situ hybridization via RNAscope®

RNAScope analysis was performed on kidneys perfused with 0.9% NaCl followed by fixation with 3% paraformaldehyde solution. The fixed tissue was dehydrated, embedded in paraffin, and cut into 5 µm sections with a microtome as described previously ^3^. Target mRNAs were hybridized and visualized using the RNAscope® Multiplex Fluorescent v2 kit (Advanced Cell Diagnostics, Hayward, CA, USA) following the manufacturer’s instructions (Wang et al., 2012). Signal detection was performed with TSA Vivid dyes 570 and 650 (Bio-Techne, Wiesbaden, Germany) and the Opal 780 fluorophore (Akoya Biosciences, Marlborough, MA). Nuclei were counterstained with DAPI included in the Multiplex Fluorescent v2 kit. Sections were mounted using ProLong™ Gold Antifade Mountant (Thermo Fisher Scientific, Waltham, MA, USA) and stored at 4 °C until further analysis. The RNAscope® probes employed are listed in Table 2.

All RNAScope® images were taken with an Axio Observer.Z1 microscope (Zeiss, Jena, Germany) using the Plan-Apochromat 20x/0.8 objective and the Colibri7 as light source.

**Table 2:** RNAscope probes used for in situ hybridization.

| RNAscope®- Probe | Cat No. | RNAscope®- Probe | Cat No. |
| --- | --- | --- | --- |
| Mm-P2ry6-C1 | 314241 | Mm-P2ry2-C3 | 406051-C3 |
| Mm-Pdgfrb-C3 | 411381-C3 | Mm-P2ry4-C2 | 406081-C2 |
| Mm-Pdgfrb-C1 | 411381 |  |  |
| Mm-P2ry1-C2 | 406061-C2 | Mm-Adgre1-C2 | 460651-C2 |

Fluorescent images were captured with the Axiocam 506 mono. Filters used were the filter set 43-Cy3 (EX BP 545/25; EM BP 605/70), filter set 50-Cy5 shift free (EX BP 640/30; EM BP 690/50), filter set 96 HE BFP (EX BP 390/40; EM BP 450/40) and filter set 115-Cy7 (EX BP 710/87; EM BP 814/91) (Zeiss). For detail fluorescent images, the Apotome.2 system (Zeiss) was used to take 10 to 15 z-stacked images, which were merged using maximum projection. Overviews were generated by stitching tiles taken at 20x magnification. Images in the same figure were taken with the same light intensities, exposure times and displayed with identical image modifications.

### FACS sorting and cell culture of murine renal fibroblasts

Murine kidneys from tamoxifen-induced PDGFR-β Cre^ERT/2^ mTmG mice were perfused with 0.9% NaCl to remove blood and 0.1 mg*mL^-1^ collagenase II-containing (Merck KGaA, Germany) in DMEM medium (PAN-Biotech GmbH, Germany). Kidneys were harvested, decapsulated, cut into small pieces and transferred to a tube with 1 mg*mL^-1^ collagenase II-containing in DMEM. Enzymatic digestion took place at 37°C at 800 rpm in a tube shaker for 60 minutes. To ensure active collagenase activity, kidney suspension was centrifuged at 3000 rpm for 3 minutes, supernatant was replaced by fresh collagenase solution three times during the incubation period. Remaining kidney fragments were dissociated by gentle pipetting using a cut 1 mL-tip. Digestion was stopped by addition of DMEM medium containing 10% fetal calf serum (FCS) (Capricorn Scientific GmbH, Germany). Cells were washed three times with PBS and resuspended for FACS sorting in PBS containing 1% FCS. Shortly before sorting, cells were transferred into a FACS tube containing a 30 µm sieve. GFP-positive cells were sorted using a BD FACSAria™ Fusion Flow Cytometer into a tube containing DMEM+10% FCS and kept on ice until cells were washed with cell culture medium (DMEM, low glucose, GlutaMax (Gibco)+15% FCS (Gibco), 1% Insulin-Transferrin-Selenium, 1%Pen/Strep+0,1% Amphotericin B) and incubated at 37°C and 5% CO_2_. Splitting of cells was done at 80-90 confluency using accutase.

### Scratch assay of FACS-sorted murine renal fibroblasts

A total of 500,000 cells were seeded into 35 mm culture dishes and grown in standard cell culture medium until reaching confluency. Cells were then serum-starved for 24 h in medium lacking fetal calf serum (FCS). Linear wounds approximately 1 mm wide were generated in the monolayer using sterile 20 μL pipette tips. After scratching, cells were washed to remove debris and dead cells, and 2 mL of fresh serum-free medium was added, supplemented with either 30 μM UDP and/or 5 μM MRS2578, or left untreated (control). Each dish contained three scratch areas. Images of the same positions were captured immediately after scratching (0 h) and at 48 h using a Zeiss Axio Observer.Z1 microscope equipped with an AxioCam 305 mono camera and a 10×/0.25 objective (Zeiss, Jena, Germany). At 48 h, cells were stained by washing with Ringer solution followed by incubation for 5 min in Ringer containing 1 μM Hoechst 33342. Migration was quantified by counting nuclei within the wound area using ZEN Intellesis software (Zeiss, Jena, Germany) using a thresholding approach. For each dish, counts from the three scratches were averaged and expressed as a fraction relative to untreated controls. Experiments were performed using at least three different FACS-sorted cell lines.

### Videomicrosopic Ca^2+^-measurements using Fura2-AM

For videomicroscopic Fura2 Ca^2+^-imaging, FACS-sorted renal fibroblasts were split at least one day prior to experiment onto glass coverslips and incubated for 30 min at 37°C with 2 µM Fura2-AM and Powerload (Roche) in Ringer solution containing (in millimoles) 5 Hepes, 145 NaCl, 5 Glucose, 0.4 KH_2_PO_4_,1.6 K_2_HPO_4_,1 MgCl_2_, 1.3 CaCl_2_. Afterwards, cells were rinsed, and glass cover slips was inserted onto a perfusion chamber and measured with the Zeiss Axio Oberver.Z1 using an Fluar 40x/1.3 oil objective (Zeiss, Jena, Germany). Cells were continuously superfused with Ringer solution containing different agonists or antagonists as indicated. Perfusion speed was 2 mL/min. Fura2 was excited at 340 and 380 nm with a LAMBDA DG-4 lamp (World Precision Instruments) using 340/40 and 387/15 BP filters and exposure times of 250 and 100 ms, respectively. Fura2 emission at 510 nm was recorded using a AxioCam 305 mono (Zeiss) and BS FT 409 and BP 510/90 filters. Sampling interval was 5 s. 340/380 emission ratio after background subtraction of cell-free region-of-interest is indicated for each cell (grey lines) within one dish. Measurements of non-responsive cells were not further analyzed as those cells were either dead or did not express necessary P2Y-receptors. For each dish, the mean basal ratios and the mean maximal ratio under agonist simulation of responsive cells was determined. Graphical summaries represent means per dish. Each experiment was at least repeated on three different days with multiple FACS-sorted primary cell lines.

### Immunofluorescence

To detect immunofluorescence signals, kidneys were perfusion-fixed with 3% paraformaldehyde and after dehydration in an ascending methanol and isopropanol series embedded in paraffin. Staining was performed on 5 μm sections. Sections were deparaffinized and blocked with 5% bovine serum albumin in phosphate-buffered saline solution and incubated with mouse α-smooth muscle actin antibody (ab7187, Abcam, Cambridge, UK) at 4°C overnight. After three washes with phosphate-buffered saline solution, sections were incubated with respective Cy3-conjugated secondary antibody (Dianova, Hamburg, Germany) and mounted with Glycergel (Agilent, Waldbronn, Germany). Overviews of one whole kidney cross-section per mouse were taken by stitching tiles with the Zeiss Axio Oberver.Z1 using an 20x/0.8 oil objective (Zeiss, Jena, Germany) equipped with an AxioCam 305 mono camera, a LAMBDA DG-4 lamp (World Precision Instruments) and filter set 43-Cy3 (EX BP 545/25; EM BP 605/70) to detect α-smooth muscle actin and filter set 38 HE (EX BP 470/40; FT 495, EM BP 525/50) to detect autofluorescence. Images in the same figure were taken with the same light intensities, exposure times and displayed with identical image modifications.

### Image Analysis

Automated image analysis (Intellesis software, Zeiss ZEN) was used to determine the cortical mRNA expression levels of PDGFR-β, P2Y6, and F4/80. Segmentation was performed by background subtraction with rolling ball method (radius 10), considering a threshold range between defined minimum intensity values (PDGFR-β: 350, P2Y6: 200, F4/80: 400) and the maximum pixel intensity (16383) with a tolerance level of 3%. No size exclusion criteria were applied during the analysis to ensure that all detected RNAscope signals were taken into account. For the detection of cell nuclei, segmentation was performed using global thresholding (DAPI intensities between 1500-16383). The area of the detected expression intensities of PDGFR-β, P2Y6, and F4/80 was normalized to the area of the detected cell nuclei. Automated analysis of cortical αSMA^+^ area normalized to EGFP-positive autofluorescence area was analyzed using a thresholding approach using the ZEN Intellesis software. Data is depicted as cortical area per mouse. Image analysis software was funded by the Deutsche Forschungsgemeinschaft (DFG, German Research Foundation) - Projektnummer 471535567.

### Statistical analyses

All data are presented as mean±SD. Data were analyzed using Origin 2024 (OriginLab Corporation, Northampton, Massachusetts, USA). In Fura2-measurements, statistical testing of agonist-signals versus baseline was performed using a paired ttest. To analyze statistical significance between different agonist stimulations, a one-way ANOVA with Tukey’s correction and mean comparisons was used. In the experimental murine fibrosis models, a Wilcoxon rank sum test was performed to analyze statistical difference except for rUUO kidneys, were contralateral and rUUO kidney were compared with a Wilcoxon signed-rank test. p values and group sizes are stated in the results section. If stated, a *post hoc* Bonferroni correction was applied for multiple testing, otherwise p ≤ 0.05 was considered statistically significant.

## NEW RESULTS

### Renal mRNA expression of G_q/11_-protein coupled P2Y receptors

To determine which murine G_q/11_-protein coupled *P2ry* (*P2ry1, P2ry2, P2ry4, P2ry6*) receptors are expressed in interstitial PDGFR-β^+^ cells of the kidney, multiplex RNA *in-situ* hybridizations on adult murine cross-sections were performed. Interstitial cells were marked using a *Pdgfrb*-probe. Note the low abundance of all P2Y-receptor mRNA compared to *Pdgfrb*, which is typical for many G_q/11_-protein coupled receptors ^31^. *P2ry* expression was seen throughout all kidney zones (Fig. 1). *P2ry1* was predominantly found in cells of the glomerulus, proximal tubular cells, urothelium and PDGFR-β^+^ interstitial cells. *P2ry2* was observed in (proximal) tubules, with increasing expression towards the medulla. *P2ry4* expression was extremely low. Few *P2ry4* signals were seen in cells of the glomerulus, (proximal) tubular cells and interstitial cells although mRNA expression seemed to be as low as one copy per cell. Interestingly, *P2ry6* was the only receptor strongly enriched in PDGFR-β^+^ fibroblasts, with additional localization in proximal tubules (1-2 mRNA copies per cell).

**Figure 1:**
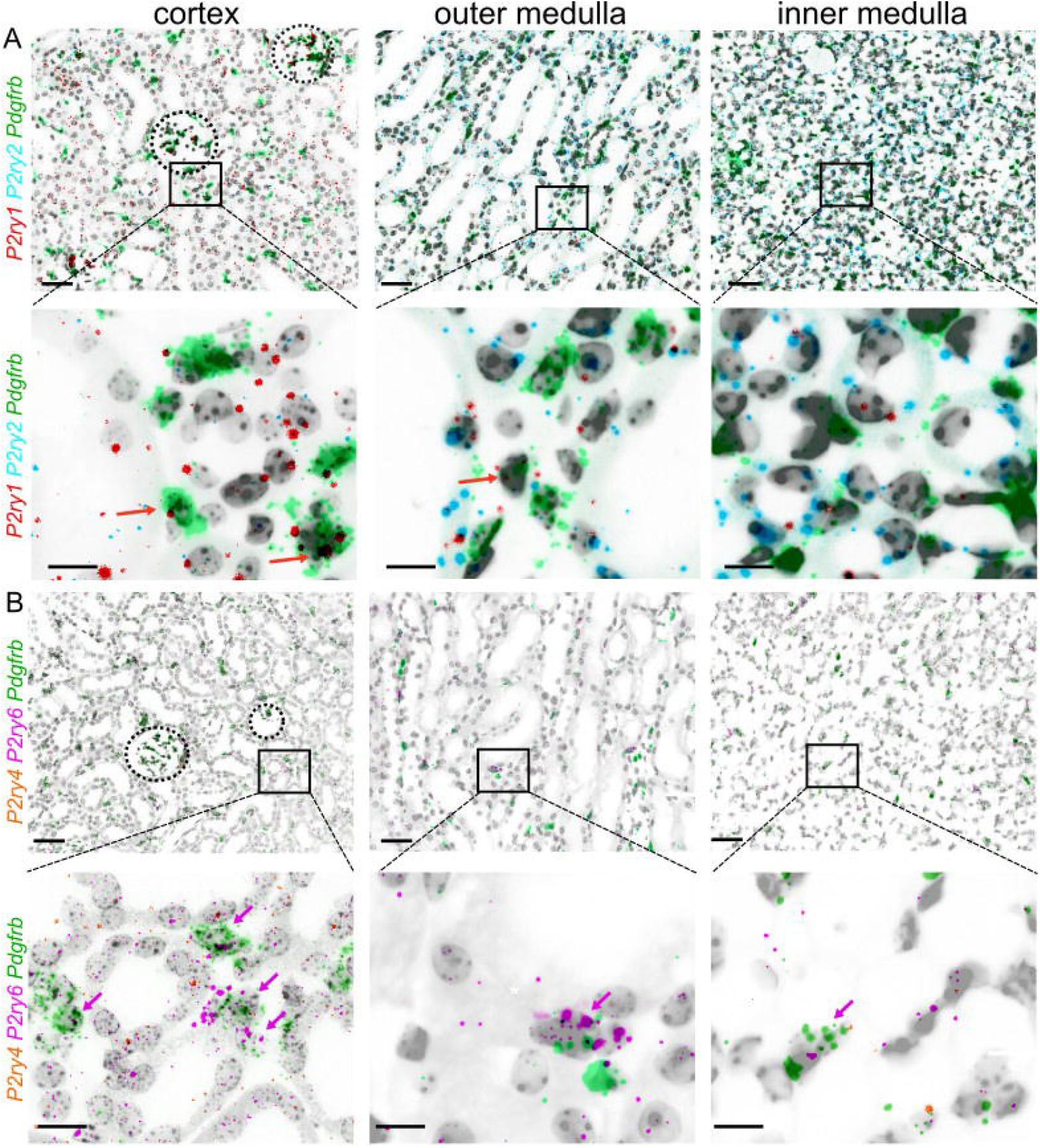
RNA expression of G_q_/11-protein coupled P2Y receptors in murine kidneys. Zonal details showing the co-expression of *P2ry1* (red),. *P2ry2* (cyan) (A), *P2ry4* (orange), *P2ry6* (pink) (B) and the interstitial fibroblast marker *Pdgfrb* (green) in transverse murine kidney sections using RNAscope. The upper panel in (A) and (B) shows details from the cortex, outer medulla and inner medulla of a control mouse (scale bar= 50 µm). The lower panel shows the respective details (scale bar= 20 µm). Colored arrows indicate double *Pdgfrb-* and respective P2yr -positive fibroblasts

### Active P2Y receptor-mediated signaling in cultured renal fibroblasts

To verify the functionality of the P2Y receptors in interstitial fibroblasts, we isolated PDGFR-β^+^ interstitial cells expressing a membranous GFP construct under the control of the *Pdgfrb*-promotor from PDGFR-β Cre^ERT/2^ mTmG mice using fluorescence-activated cell sorting (FACS). After cultivation, we verified expression of the G_q/11_-protein coupled P2Y receptors using mRNA *in-situ* hybridization (Supplementary Fig. 1). Subsequently, fibroblasts were loaded with the Ca^2+^-indicator Fura-2 and active G_q/11_-protein signaling was investigated using videomicroscopic imaging. Cells responded to superfusion with the nucleotides ATP, ADP, UTP or UDP with transiently increased oscillating Ca^2+^-signals indicative of intracellular store release (Figure 2). P2X involvement was excluded by removal of extracellular Ca^2+^, which did not affect ATP-mediated cytosolic Ca^2+^-signaling (Supplementary Fig 2A). To identify active P2Y isoforms, we applied selective antagonists where available. We observed that ADP-mediated signaling could be significantly reduced, but not completely suppressed, by specific inhibition of the ADP-sensitive P2Y_1_ receptor using MRS2179 (Fig. 2B). Similarly, UDP-mediated responses were markedly attenuated by MRS2578, an irreversible covalent antagonist of P2Y_6_ that inactivates the receptor by modifying a critical cysteine residue, leading to internalization and degradation ^32,33^ (Fig. 2C). Notably, neither MRS2179 nor MRS2578 caused nonspecific inhibition of Ca^2+^ responses to other nucleotides, whereas the broad-spectrum P2 receptor antagonist suramin suppressed all nucleotide-induced Ca^2+^ transients (Supplementary Fig. 2).

**Figure 2:**
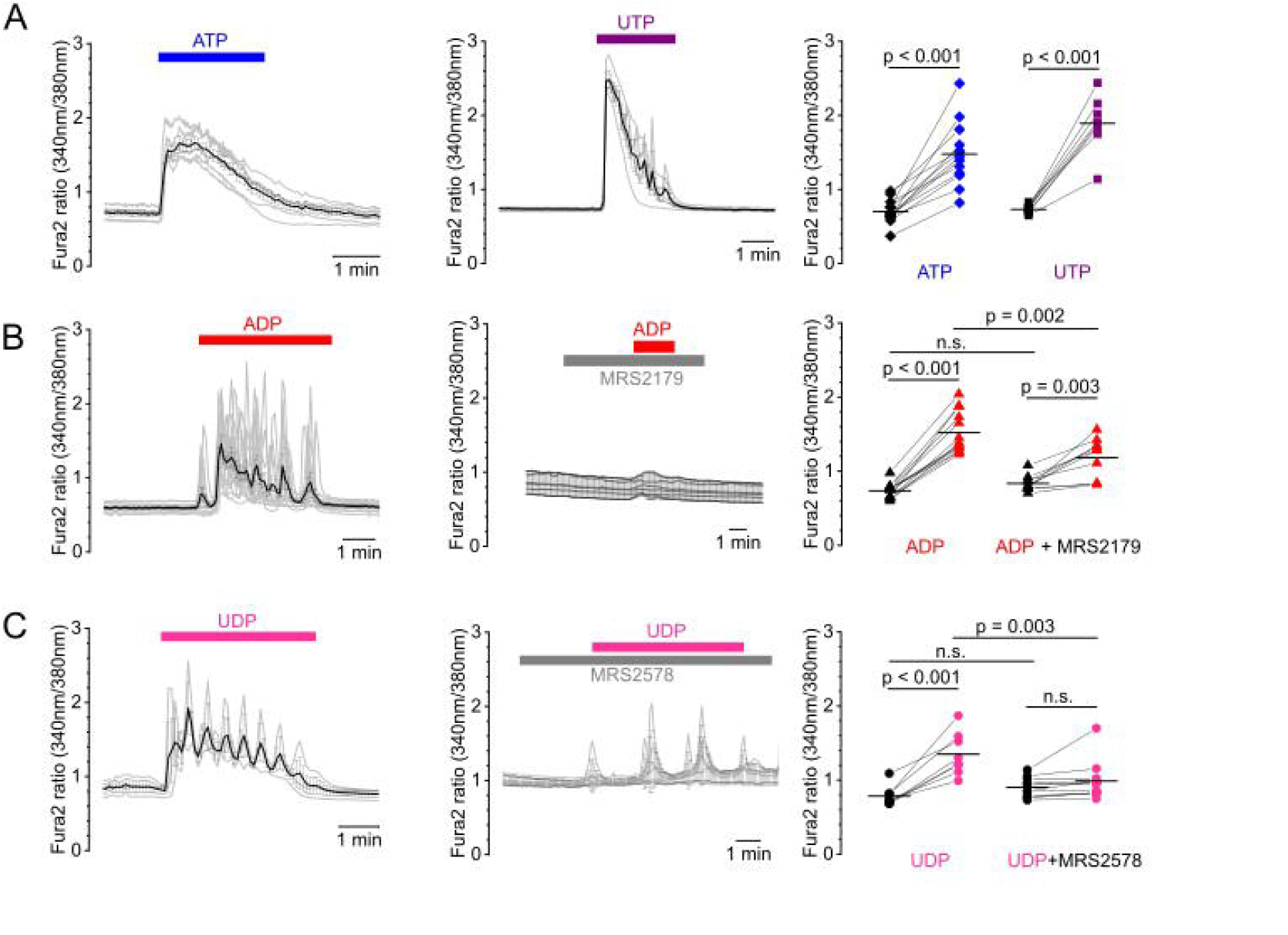
Active P2Y receptor-mediated signaling in cultured renal fibroblasts. (A) FACS-sorted, cultivated murine renal fibroblasts were loaded with the Ca**2**^**+**^-indicator Fura2 and superfused with agonists and antagonists of P2Y receptors. (A) Representative trace of single cells (grey lines) superfused with 100 µM ATP (left) or U TP (middle). Black lines represent the mean traces per dish. Summary on the right indicates basal (black) and respective maximal Ca^**2**+^-transients (colored symbols) under activation. Each connected dot represents the mean of one dish. (B, C) Left panel shows a representative trace of cells superfused with 100 µM ADP (B) or UDP (C). Middle panel depicts a representative trace in the presence of 5 µM of the specific P2Y_**1**_ inhibitor MRS2179 (B) or 5 µM of the specific P2Y_**6**_ inhibitor MRS2578 (C). Summary on the right indicates basal (black) and respective maximal Ca^**2+**^-transients (colored symbols) of the respective measurements. Each condition was tested at least with three different cell lines.

### P2Y_6_ receptors are upregulated in renal fibrosis models

To analyze the role of G_q/11_-protein coupled P2Y receptors in fibrosis, we re-examined cDNA (generated in previous study^3^ from whole kidney lysates of different experimental kidney fibrosis models including adenine nephropathy (adenine) and unilateral ureteral obstruction (UUO) using quantitative PCR (Supplementary Fig. 3). *P2ry1* mRNA normalized to the housekeeping gene *actb* was significantly downregulated from 1.00±0.28 to 0.69±0.45 the UUO model (p=0.018), but not the in adenine-induced nephropathy (p=0.268), while *P2ry2* mRNA was significantly downregulated from 1.00±0.3 in the UUO model (p=0.008) but significantly upregulated in the adenine-induced nephropathy from 1.00±0.46 to 1.64±0.22 (p=0.042) (Supplementary Fig. 3). We could not reproducibly amplify *P2ry4* mRNA probably due its low expression as evidenced by the RNA *in situ* hybridization experiments. Normalized *P2ry6* was increased in both fibrosis models from 1.00±0.53 to 1.85±1.26 in the UUO model (p=0.032) and from 1.00±0.70 to 3.83±1.39 in the adenine model (p=0.002). These results are indicative for distinctively different regulation of G_q/11_PCR P2Y-receptor isoforms in renal disease progression with P2Y_6_ upregulation being a common entity. To analyze the localization of renal P2Y_6_ receptors under fibrotic conditions, we conducted a reversible unilateral ureter obstruction (rUUO) for five days whereafter the fibrosis progression was continued for two weeks. Similarly, adenine-induced nephropathy was induced by a high adenine diet for three weeks whereafter kidneys were harvested and analyzed using RNA *in situ* hybridization with specific *P2ry6* and *Pdgfrb*-probes (Fig. 3). Automated quantitative image analysis of cortical *P2ry6* signals revealed that, regardless of the disease model, the 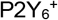 area was upregulated from 3.3±0.5% to 5.9±1.5% in the rUUO model (p=0.042) where the contralateral kidney served as internal control, and from 4.5±0.9% to 11.3±4.3 in the adenine model (p<0.001) (Fig. 3D). Interestingly, *P2ry6*^+^ signal was strongly enriched in the interstitial space but not exclusively co-localized with *Pdgfrb*. Besides *Pdgfrb*^+^-co-hybridization, we observed a strong co-localization with the macrophage marker F4/80 (*Adgre1*) that also increased in the cortical areas from 1.9±0.6% to 4.1±1.3% in the rUUO model (p=0.032) and from 1.0±0.3% to 7.6±3.4% in the adenine model (p<0.001). In contrast, *Pdgfrb*^+^ area did not significantly change in the disease models.

**Figure 3:**
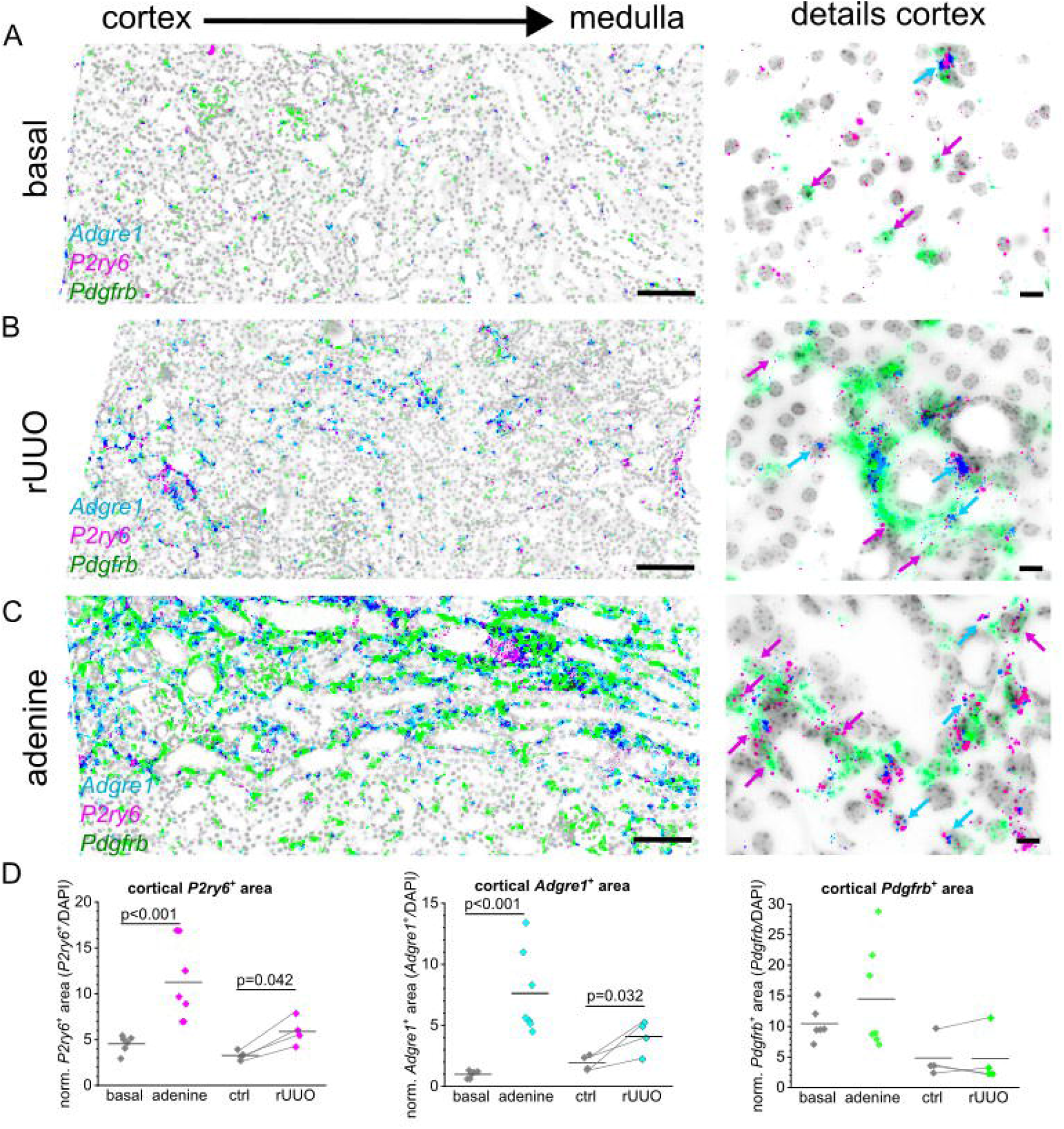
*P2ry6* is regulated in renal fibrosis models. Representative pictures on the left show the co-expression of *P2ry6* (pink), the macrophage marker *Adgre1* (cyan) and the interstitial fibroblast marker *Pdgfrb* (green) in transverse murine kidney sections using RNAscope (scale bar= 100 µm), while close-ups on the right present cortical details (scale bar= 20 µm) of a control mouse (A). Representative fibrotic kidney sections from either a mouse subjected to reversible unilateral ureter obstruction (rUUO), or adenine-induced nephropathy are depicted in (B) or (C) respectively. (D) Automated image analysis of cortical *P2ry6+* (left), *Adgre1* + (middle) or *Pdgfrb+* (right) area normalized to total analyzed area. Each dot represents one mouse. In rUUO mice, contralateral kidneys (ctrl) were used as control and respective damaged kidney are linked by a line.

### Pharmacological inhibition of P2Y_6_ receptors in fibrosis progression

To assess the role of P2Y_6_ in the development of renal fibrosis, we inhibited P2Y_6_ receptors using MRS2578 in adenine-induced nephropathy. Fibrosis progression was assessed by immunofluorescent staining of α-smooth muscle actin (αSMA) on transverse murine kidney slices. The percentage of αSMA^+^ area was determined automatically using a thresholding approach (Fig. 4). Adenine diet resulted in statistically significant increase of cortical αSMA^+^ area compared to control kidneys (p=0.002). Pharmacological inhibition of P2Y_6_ using MRS2578 significantly attenuated fibrosis progression as estimated by αSMA immunostaining from 1.00±0.31 to 0.53±0.16 (p=0.004) in the adenine-induced model, with the mean value for vascular αSMA in healthy control kidneys being 0.29 ± 0.14. Additionally, the mRNA expression of markers of fibrosis was examined using qPCR. As depicted in Fig. 4D, collagen I (*Col1a1*) mRNA levels were significantly decreased in MRS2578 treated animals compared to animals receiving vehicle injections and adenine enriched food. P2Y_6_ (*P2ry6*) mRNA levels were also examined. They were upregulated in the adenine-fed animals, but not significantly affected by MRS2578.

**Figure 4:**
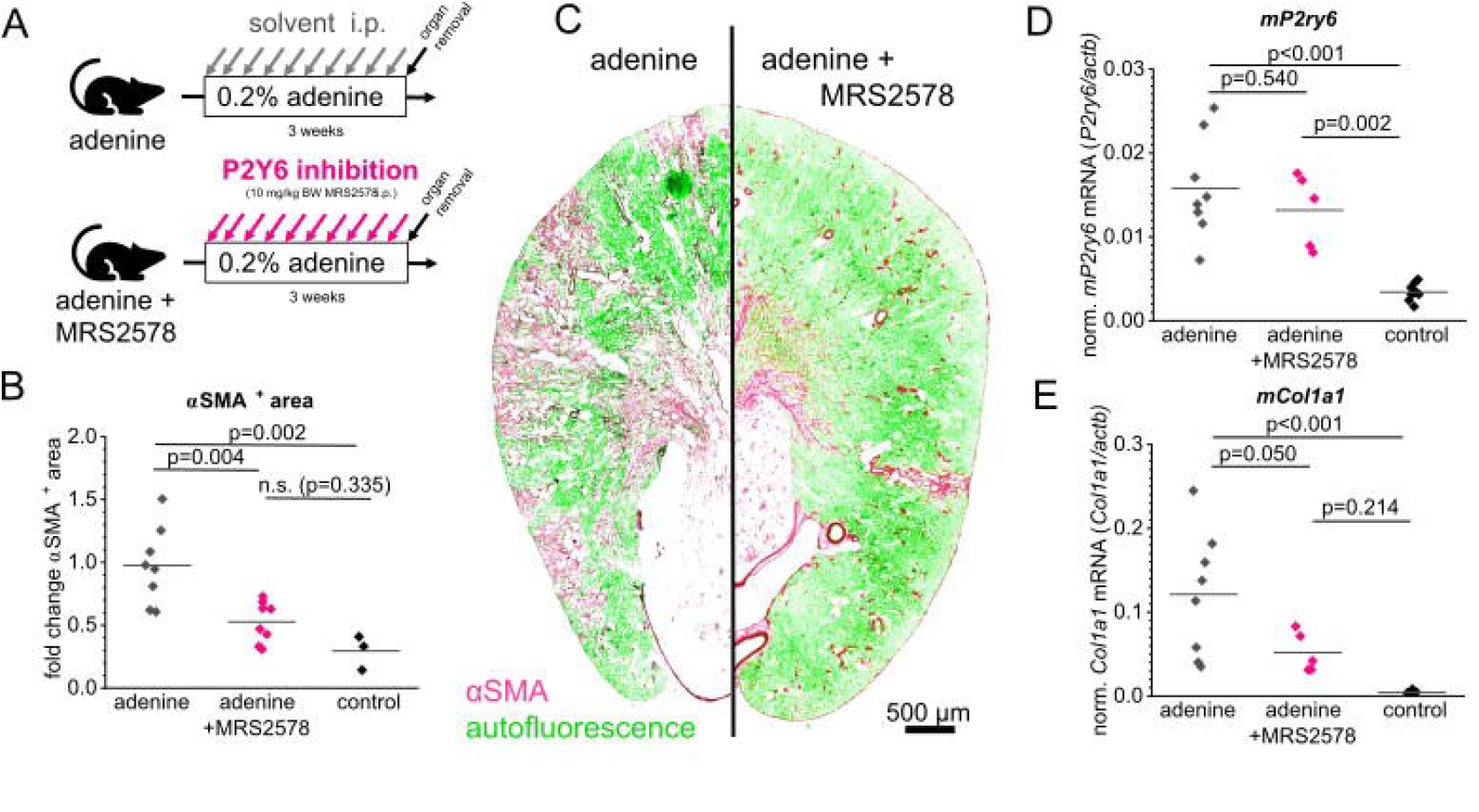
Inhibition of P2Y6 signaling in mice attenuates fibrosis progression. (A) To induce a renal fibrosis, male mice were fed a high adenine diet (0.2% adenine) for three weeks. During this period, one group of animals was injected i.p. with 10 mg/kg BW of the specific P2Y_**6**_ inhibitor MRS2578 (marked “adenine + MRS2578”). The group marked with “adenine” received a diet enriched with adenine and injections of the vehicle. The extent of fibrosis was automatically analyzed using transverse murine kidney sections of each mouse that were positive for aSMA staining (B). (C) Representative kidney sections of both groups showing aSMA staining in pink, while green color indicates autofluorescence. (D, E) Quantitative real time analysis of *P2ry6* mRNA levels (D) normalized to actb as well as collagen I *(Co/1a1)* mRNA levels (E). Abbreviations: aSMA = a smooth muscle actin.

### P2Y_6_ receptor activation promotes migration of fibroblasts

Given the expression of P2Y_6_ in interstitial cells and macrophages, we wondered what effect P2Y_6_ activation has on these cells. Macrophages are known to be activated in the presence of the P2Y_6_ agonist UDP leading to a pro-inflammatory, chemokine-releasing phenotype ^34,35^. Given that UDP acts as a danger associated molecular pattern in the interstitium, we speculated that PDGR-β^+^ fibroblasts might be drawn to the site of UDP release. Therefore, we examined migration of serum-starved cultured FACS-sorted PDGFR-β^+^ fibroblasts in a wound healing assay, where a confluent monolayer of fibroblasts was scratched with a pipette tip and photographed immediately and 48h after the scratch formation. Cells at the 48h time point were additionally stained using the nuclear marker HOE33342 and migration of cells into the scratch was automatically analyzed by counting nuclei of invading cells (Fig. 5). Migration was assessed by normalizing to invading cells in control conditions. In the presence of 30 µM UDP, 14.6±13.1% more cells migrated into the scratch compared to control cells (p=0.003) while the addition of 5 µM MRS2578 abolished the effect of UDP (-7.0±15.1%, p=0.303). The addition of MRS2578 alone also significantly decreased migration by -20.3±6.3% (p=0.002).

**Figure 5:**
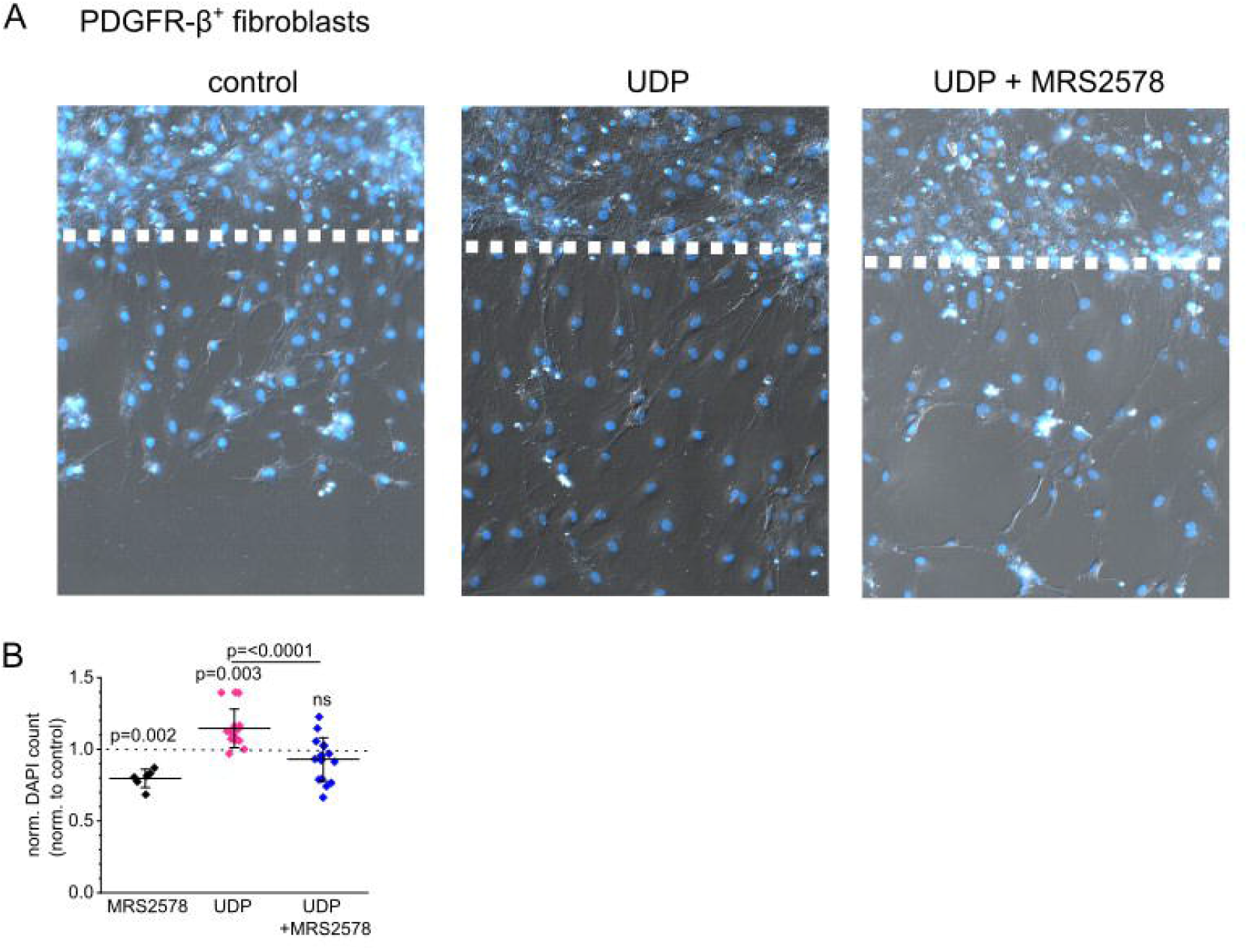
UDP promotes cell migration of cultured renal fibroblasts via P2Y_6_ activation. (A) Representative pictures of cultured FACS sorted murine PDGFR-β-positive renal fibroblasts 48h after wounding and subsequent stimulation with either 30 µM UDP or 30 µM UDP + 5 µM of the P2Y6 inhibitor MRS2578. Nuclei were counterstained using HOE33342 indicated in blue. (B) Migration was evaluated by automatic counting of migrated nuclei per area and is depicted as a percentage compared with the control (stippled line). Values are expressed as means + STD from eight independent experiments. Abbreviations: FACS = fluorescence-activated cell sorting, PDGFR-β= platelet-derived growth factorβ. UDP = Uridine Diphosphate

## DISCUSSION

Renal fibrosis is a central pathological feature of chronic kidney disease (CKD), ultimately leading to irreversible loss of renal function. More than 850 million people worldwide are affected by CKD, making it the 12^th^ leading cause of death globally, and it is projected to become the 5^th^ leading cause by 2040 ^36^. CKD is characterized by excessive accumulation of extracellular matrix (ECM), primarily produced by activated myofibroblasts, which largely originate from PDGFR-β^+^ interstitial fibroblasts ^4,6,104^. Understanding the molecular mechanisms driving fibroblast activation is crucial for developing targeted anti-fibrotic therapies.

In this study, we investigated the role of G_q/11_-protein coupled P2Y receptors in renal interstitial fibroblasts and their contribution to experimental kidney fibrosis. Among all P2Y receptors analyzed, only P2Y_1_ and P2Y_6_ were enriched in renal interstitial cells, with P2Y_6_ being selectively upregulated in experimental fibrosis models. Strikingly, pharmacological interference in the P2Y_6_ signaling during fibrosis progression in mice significantly attenuated fibrosis outcome in mice. These findings expand the current understanding of purinergic signaling in renal (patho)physiology and highlight its relevance in fibroblast regulation.

While P2 receptor expression in renal epithelial and vascular cells is well documented, their role in interstitial fibroblasts—major drivers of kidney fibrosis—has remained largely unexplored. Due to the low mRNA copy number as well as low protein abundance of many (G-protein coupled) receptors, we used the highly specific mRNA hybridization technique RNAScope to explore localization of the G_q/11_-protein coupled P2Y receptors P2Y_1_ (*P2ry1*), P2Y_2_ (*P2ry2*), P2Y_4_ (*P2ry4*), and P2Y_6_ (*P2ry6*) in renal interstitial fibroblasts that we labeled with a PDGFR-β (*Pdgfrb*) probe (Fig. 1). P2Y_1_ was detected in glomerular cells, proximal tubular cells, urothelium and PDGFR-β^+^ interstitial cells. Consistent with the literature reporting P2Y_2_ activity in (proximal) tubular cells, mesangial cells and arterioles, we observed P2Y_2_ localization in epithelial cells, with increasing expression towards the medulla ^15,20^. P2Y_4_ expression was minimal, while P2Y_6_ was the only receptor strongly enriched in PDGFR-β^+^ fibroblasts, with additional localization in proximal tubules.

Functional assays confirmed receptor activity: superfusion of FACS-sorted cultured renal fibroblasts with different nucleotides induced Ca^2+^-transients (Fig. 2). Specific pharmacological interference of either P2Y_1_ using MRS2179 or P2Y_6_ using MRS2578 did significantly reduce ADP- or UDP-mediated Ca^2+^-signals although Ca^2+^-signals were not completely abolished supporting receptor specificity despite partial cross-activation at high nucleotide concentrations ^37^. Of note, we did not detect unspecific blockage of other nucleotide signals using MRS2179 or MRS2578 verifying the specificity of the used compounds (Supplementary Fig. 2). The distinct expression profile of the different P2Y receptors was also evident in RNAScope experiments using the same FACS-sorted cell lines (Supplementary Fig. 1) where P2Y_4_ expression was low and P2Y_6_, P2Y_1_ and P2Y_2_ expression was abundant suggesting that although the components of the nucleotide signaling pathway are redundantly expressed, each P2Y seems to have a distinct role in cell regulation.

Importantly, P2Y_6_ expression was markedly upregulated in two murine fibrosis models — adenine-induced nephropathy and unilateral ureteral obstruction — while other P2Y receptors were downregulated. RNAscope revealed P2Y_6_ localization not only in PDGFR-β^+^ fibroblasts but also in F4/80^+^ macrophages, suggesting a dual role in fibroblast activation and immune cell recruitment (Fig. 3).

We wondered how these findings in mice translate to a human kidney injury and attempted immunofluorescent labeling of P2Y_6_ in murine and human tissue samples. Attempts to validate P2Y_6_ protein expression by immunofluorescence were inconclusive, likely due to low receptor abundance. However, searches in commonly available human databases verified our own observations since P2Y_6_ upregulation in fibroblasts was also observed in an single-cell transcriptomics approach with acute kidney injury samples ^38^ as well as in single cells RNA sequencing data from the KPMP kidney tissue atlas (https://atlas.kpmp.org) indicating a common cross-species phenomena in mice and men.

Given the expression of P2Y_6_ in the renal interstitium, the question arose whether inhibition of P2Y_6_-signaling using the specific inhibitor MRS2578 might affect fibrosis progression in experimental fibrosis models (Fig. 4). The most striking observation of our study was that pharmacological inhibition of P2Y_6_ with MRS2578 using i.p. injections three times a week significantly reduced fibrosis in adenine-induced nephropathy, as evidenced by decreased αSMA^+^ myofibroblast area and lower expression of fibrotic markers such as collagen I (*Col1a1*). These findings underscore the therapeutic potential of targeting P2Y_6_ in CKD.

In the face of the multifactorial modes of action it is not surprising that dysregulation of nucleotide signaling pathway is involved in numerous diseases like fibrosis models, polycystic kidney disease, diabetic nephropathy, ischemia, nephritis, inflammation and hypertension. Moreover, abrogation of specific purinergic players like pannexin 1, P2X_7_, P2Y_12_ or P2Y_2_ have been shown to be beneficial in the progression of the respective disease ^16,20^. However, clinically available drugs only include the P2Y_12_ antagonist that inhibits thrombosis, while numerous completed phase 2 trials on P2X_7_ antagonists did not show the expected benefit ^24^.

It is intriguing to speculate how interstitial fibroblasts, which predominantly express diphosphate-sensitive receptors P2Y_1_ and P2Y_6_, are activated under physiological conditions. While ATP release is well characterized — occurring via exocytosis, ATP-permeable channels such as connexin 43 (*Cx43*) and pannexin 1 (*Panx1*), or passively from dying cells — the mechanisms governing UTP or UDP release are less understood, though release from necrotic cells appears likely ^16,17^. Studies in rat intestinal epithelial cells revealed negligible basal extracellular nucleotide levels, yet mechanical injury triggered rapid release of ATP and UDP, but not ADP or UTP ^39^, suggesting distinct nucleotide-release dynamics.

What are the consequences of UDP release for effector cells? Our data indicate that UDP enhances fibroblast migration in wound-healing assays (Fig. 5). This suggests that PDGFR-β^+^ fibroblasts may migrate toward sites of UDP release. Conversely, macrophages exhibited reduced migration under the same conditions. Previous studies have shown that macrophages are attracted by UDP but subsequently activate and differentiate ^34^. It is intriguing to speculate, that P2Y_6_ signaling may also influence macrophage infiltration indirectly by modulating pericyte contraction and detachment, leading to heterogeneous capillary constriction, focal hypoperfusion, and localized barrier disruption. These changes could create “hot spots” of damage that facilitate macrophage entry. As pericytes transition into myofibroblasts during chronic injury, loss of pericyte coverage and deposition of stiff extracellular matrix may permanently alter capillary geometry, generating hypoxic niches that favor pro-fibrotic macrophage phenotypes. Future studies are needed to evaluate the contribution of PDGFR-β^+^ and macrophages to this phenomenon.

Interestingly, Bar et al., observed that P2Y_6_-null mice are viable and phenotypically indistinguishable from wild-type mice in terms of growth and fertility. Yet, they exhibit impaired UDP responses in macrophages, endothelial cells, and vascular smooth muscle ^34^. The same group also implicated P2Y_6_ to be a therapeutic target to regulate cardiac hypertrophy since the P2Y_6_ gene knockout is associated with a macrocardia phenotype and amplified pathological cardiac hypertrophy in mice ^40^. Beyond renal and cardiovascular disease, recent evidence suggests P2Y_6_ as potential immunotherapy target since increased production of UDP attracts immunosuppressive macrophages through its receptor P2Y_6_ while pharmacological interference of immunosuppressive macrophages by MRS2578 promoted responsiveness to immunotherapies in otherwise resistant pancreatic ductal adenocarcinoma and melanoma models ^41^.

In summary, this study demonstrates that P2Y_6_ is expressed under control conditions in renal interstitial fibroblasts and, upon activation, promotes cell migration. In CKD, P2Y_6_ is additionally expressed by infiltrating macrophages. Pharmacological blockade with MRS2578 markedly reduced fibrotic lesions, highlighting its therapeutic potential. Our study is limited by the lack of P2Y_6_ protein level verification since a specific antibody to verify P2Y_6_ protein expression is lacking. Secondly, our study is based on murine models, although single-cell RNA sequencing data suggest conservation in humans. Nonetheless, functional validation in human tissue or organoid models is needed. Third, the pharmacological inhibitor MRS2578, while selective, may have off-target effects, and its pharmacokinetics and safety profile remain insufficiently characterized for clinical translation which should be addressed in future studies. Together with prior studies highlighting P2Y_6_ inhibition as beneficial and the absence of major phenotypic abnormalities in global P2Y_6_ knockout mice, these results position P2Y_6_ as a promising candidate for further investigation into fibrotic signaling and targeted therapy development for CKD.

### Outlook

The interplay between interstitial fibroblasts and the purinergic signaling pathway represents a promising starting point for the development of new antifibrotic therapies that might target interstitial fibroblasts and macrophages at the same time. A deeper understanding of these molecular mechanisms could make a decisive contribution to slowing down or even stopping the progression of chronic kidney disease.

**The discussion is in part based upon data generated by the Kidney Precision Medicine Project. Accessed December 8th, 2025. https://www.kpmp.org.**

## ACKNOWLEDGEMENTS

The authors would like to thank Ines Tegtmeier, Linh Minh Tran and Justina Rötsch for their expert technical assistance. This work was funded by **the Deutsche Forschungsgemeinschaft (DFG, German Research Foundation), project number 509149993, TRR 374**.

The discussion is in part based upon data generated by the Kidney Precision Medicine Project. Accessed December 8th, 2025. https://www.kpmp.org. The Kidney Precision Medicine Project (KPMP) is supported by the National Institute of Diabetes and Digestive and Kidney Diseases (NIDDK) through the following grants: U01DK133081, U01DK133091, U01DK133092, U01DK133093, U01DK133095, U01DK133097, U01DK114866, U01DK114908, U01DK133090, U01DK133113, U01DK133766, U01DK133768, U01DK114907, U01DK114920, U01DK114923, U01DK114933, U24DK114886, UH3DK114926, UH3DK114861, UH3DK114915, and UH3DK114937. We gratefully acknowledge the essential contributions of our patient participants and the support of the American public through their tax dollars.

## Notes

### Competing Interest Statement

The authors have declared no competing interest.

