## Supplementary figures and images for "Inhibition of (interstitial) P2Y_6_ receptors attenuates renal fibrosis progression"

### Supplementary Fig. 1

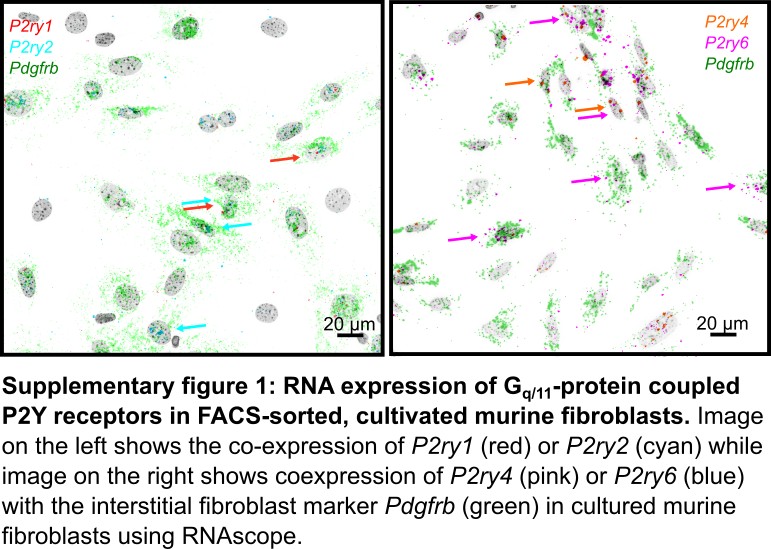

### Supplementary Fig. 2

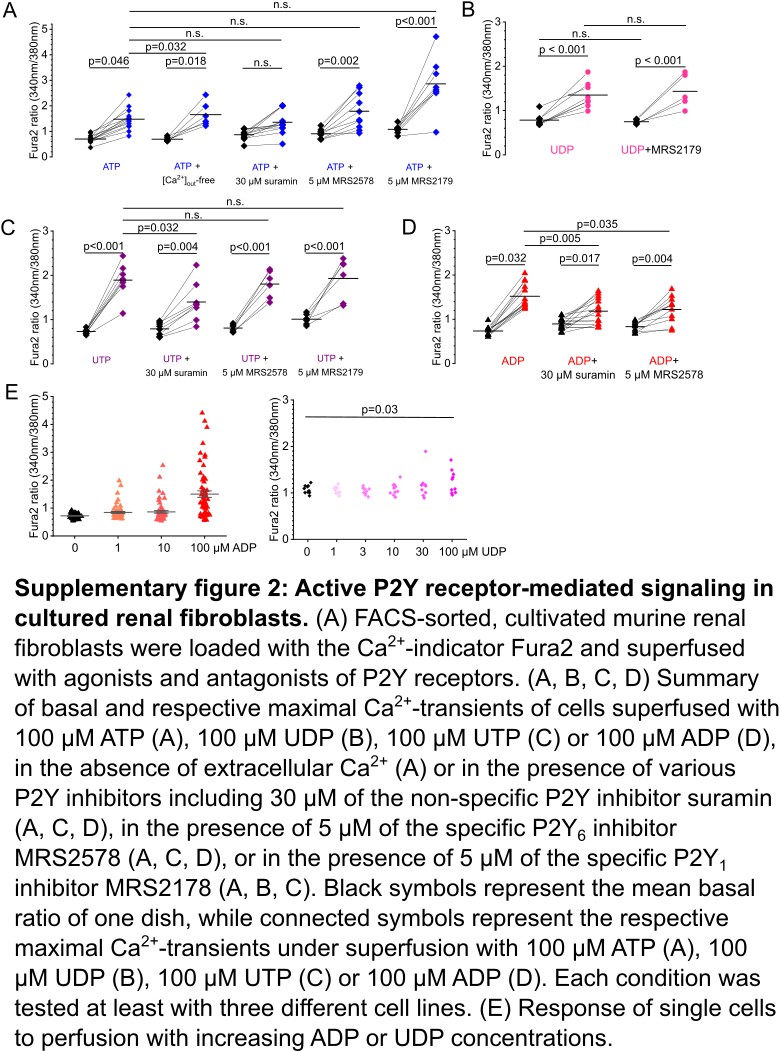

### Supplementary Fig. 3

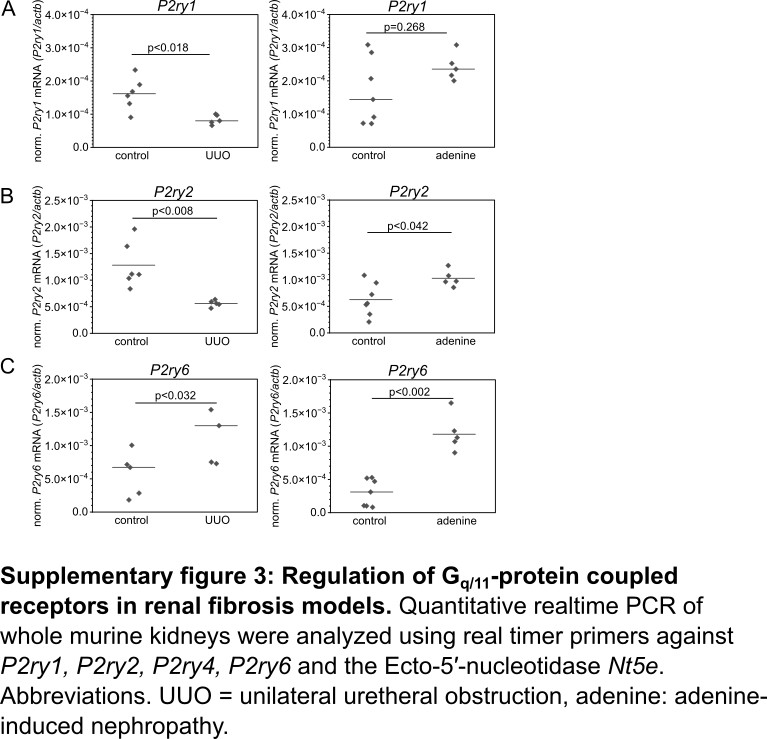
